# Methylome analysis reveals dysregulated developmental and viral pathways in breast cancer

**DOI:** 10.1101/034322

**Authors:** Mohammed OE Abdallah, Ubai K Algizouli, Maram Abbas Suliman, Rawya Abdulaziz Abdulrahman, Mahmoud Koko, Ghimja Fessahaye, Jamal Haleem Shakir, Ahmed H. Fahal, Ahmed M Elhassan, Muntaser E Ibrahim, Hiba S Mohamed

**Affiliations:** Institute of Endemic Disease, University of Khartoum, P. O. Box 102. Khartoum Sudan; Khartoum teaching hospital. Khartoum. Sudan; Faculty of Medicine. University of Khartoum. Khartoum. Sudan

**Keywords:** Methylome, Breast Cancer, Epigenetics, DNA Methylation, HM450, Epigenome Reference

## Abstract

**Background:** Breast cancer (BC) ranks among the most common cancers in Sudan and worldwide with hefty toll on female health and human resources. Recent studies have uncovered a common BC signature characterized by low frequency of oncogenic mutations and high frequency of epigenetic silencing of major BC tumor suppressor genes. Therefore, we conducted a genome-wide methylome study to characterize aberrant DNA methylation in breast cancer.

**Results:** Differential methylation analysis between primary tumor samples and normal samples from healthy adjacent tissues yielded 20188 differentially methylated positions (DMPs), which is further divided into 13633 hypermethylated sites corresponding to 5339 genes and 6555 hypomethylated sites corresponding to 2811 genes. Moreover, bioinformatics analysis revealed epigenetic dysregulation of major developmental pathways including hippo signaling pathway. We also uncovered many clues to a possible role for EBV infection in BC

**Conclusion:** Our results clearly show the utility of epigenetic assays in interrogating breast cancer tumorigenesis, and pinpointing specific developmental and viral pathways dysregulation that might serve as potential biomarkers or targets for therapeutic interventions.

## Background

Breast cancer (BC) is the most common cancer among females in Sudan [1–3], and is still a leading cause of high morbidity and mortality across the world. According to a recent report from the national cancer registry[2], BC had an incidence rate of 25.1 per 100.000, more than twice the incidence rate of the second commonest cancer. Furthermore, Sudanese BC patients tend to present at young age, at late stage, and with advanced disease compared to their counterparts in other countries [4]. Another study [5] reported a young age of presentation for locally advanced BC. Therefore, there is an urgent need for serious epidemiologic and molecular studies in order to trace the underlying mechanisms behind BC, and for developing better early detection methods as well as a nationwide educational effort to tackle this ravaging disease.

Epigenetics has emerged as a new, rapidly growing field in biology, with significant implications for cancer research. Epigenetic modifications include DNA methylation, and histone modifications, although they both do not alter DNA sequence per se, they influence chromatin remodeling and thus offer a dynamic and flexible way of controlling gene expression.

DNA methylation of cytosine residues occurs predominantly at CpG sites, and is mediated by three DNA methyltransferases (DNMTs). DNMT1, which maintains DNA methylation during cell replication, and a pair of DNMT3s – DNMT3a and DNMT3b – which is responsible for de novo DNA methylation. Epigenetic reprogramming through genome-wide alteration of DNA methylation (methylome) is critical for control of development and differentiation in normal cells and tissues, however, faulty epigenetic reprogramming, as in aberrant DNA methylation, can be a major driver of multiple types of cancer including BC [6, 7]

Methylome analysis has proved to be very pertinent to the study of the different aspects of cancer tumorigenesis. The vast majority of methylation changes occurs in a tissue-specific manner [8], which makes methylome profiling a very sensitive and specific method for delineating dysregulated epigenetic pathways at the tissue level, as in cancer, which usually arises from a single tissue. Moreover, DNA methylation is a stable epigenetic mark that is ideal for development of biomarker assays, which can offer a rapid, cost effective, a and minimally invasive diagnostic/prognostic tests [9, 10]. Additionally, methylome analysis has been effectively used in tumor subtype classification [11–15]. Furthermore, genome-wide methylome assays have also proved to be very useful in detecting and profiling viral epigenetic signature in cancer [16–18].

The aim of the present study was to investigate genome-wide DNA methylation profile of breast cancer in Sudanese patients utilizing Illumina Infinium HumanMethylation450 BeadChips (HM450) methylation assay. This array-based assay is widely used in epigenetics studies, and is a reliable, cost effective, high throughput method. We conducted methylome analysis comparing primary BC tissue samples against normal samples from adjacent healthy tissues. The results of this study provide a valuable insight into the epigenetics of BC in Sudanese patients.

## Results

### Genome-wide DNA Differential methylation Analysis

Each of three approaches – listed in Materials and Methods-produced a list of differentially methylated sites: Limma, 39940; Wilcoxon, 34099; Nimbl, 22251 (0.2 median beta value difference, Benjamini-Hochberg adjusted p-value ≤0.05). Here we only report the results for final set obtained from Nimbl-compare module, which represents the intersection of the three methods. The final set consisted of 20188 differentially methylated CpG sites, which is further divided into 13633 hypermethylated sites corresponding to 5339 genes and 6555 hypomethylated sites corresponding to 2811 genes. Nimbl unique approach ensured detection of differentially methylated positions (DMPs) that have the largest effect size as illustrated in **Fig 1**. A volcano plot showing the demarcation of differentially methylated sites by both statistical significance and effect size is shown in **Fig 1.B** Hierarchical clustering of the top 250 differentially methylated sites sorted by F value (low intragroup variability and higher intergroup variability) is shown in **Fig 2**. The resulting heatmap and dendogram showed clear separation of tumor samples from normal samples

**Figure 1:**
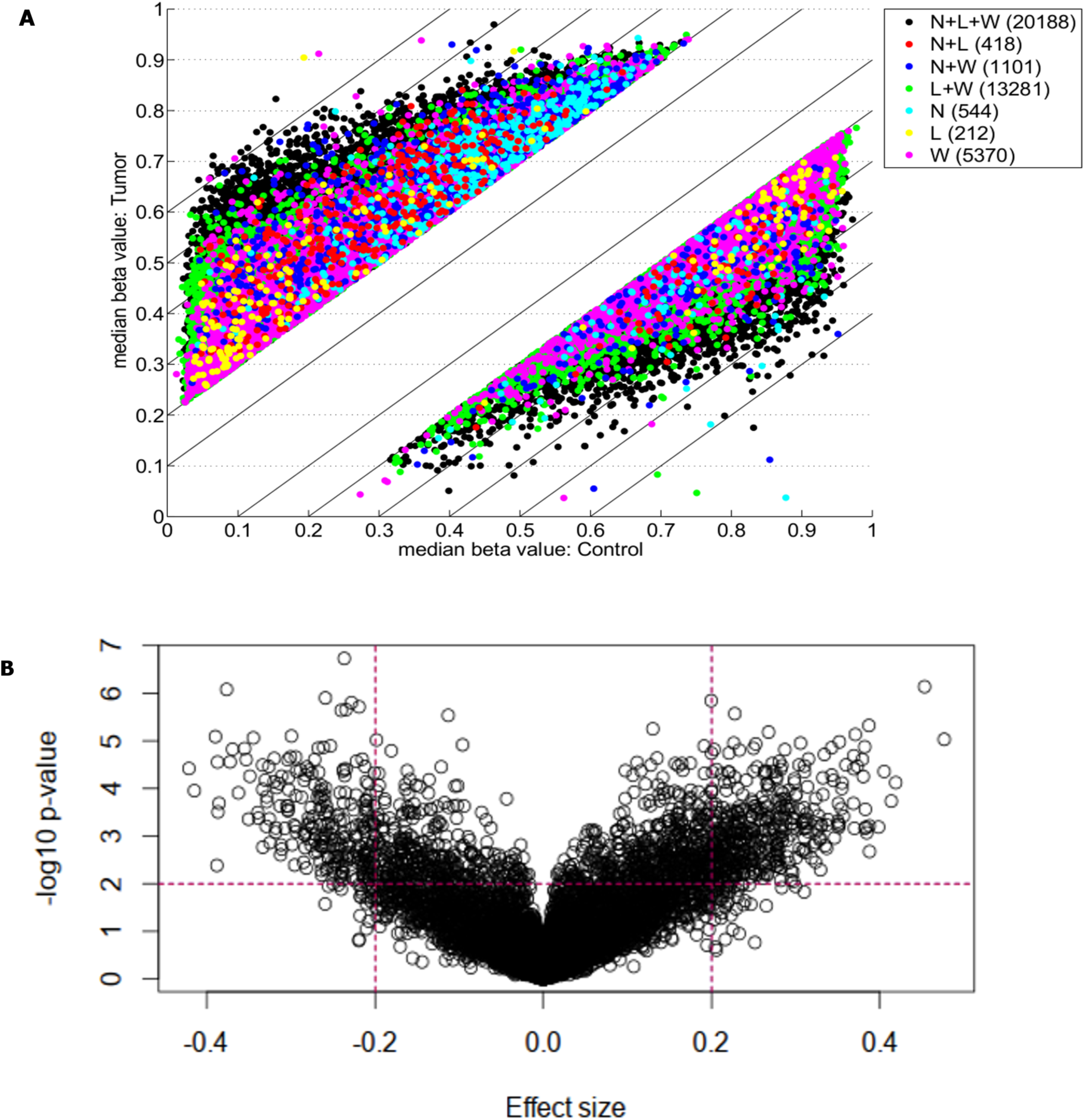
Genome-wide DNA Differential methylation Analysis of study samples. (A) shows differentially methylated CpG sites (defined as median beta value difference equal to or more than 0.2) identified using three methods: Limma (L; 34099 sites), Wilcoxon (W; 39940 sites), and Nimbl, (N; 22251 sites). The color code shows sites identified by each method alone and in combination. A final set which represents the intersection of three approaches (L + W + N; black dots) consisted of 20188 sites was obtained by Nimble-compare module and used for analysis in this study. (B) A volcano plot showing the demarcation of differentially methylated sites by both statistical significance and effect size. The sites targeted in this study are those with high effect size (median beta value difference equal to or more than 0.2) and low p-value (equal to or more than 0.01, shown as -log10). The dotted lines show these cut-offs. Targeted sites for analysis are those in outer upper rectangular area of the plot.

**Figure 2:**
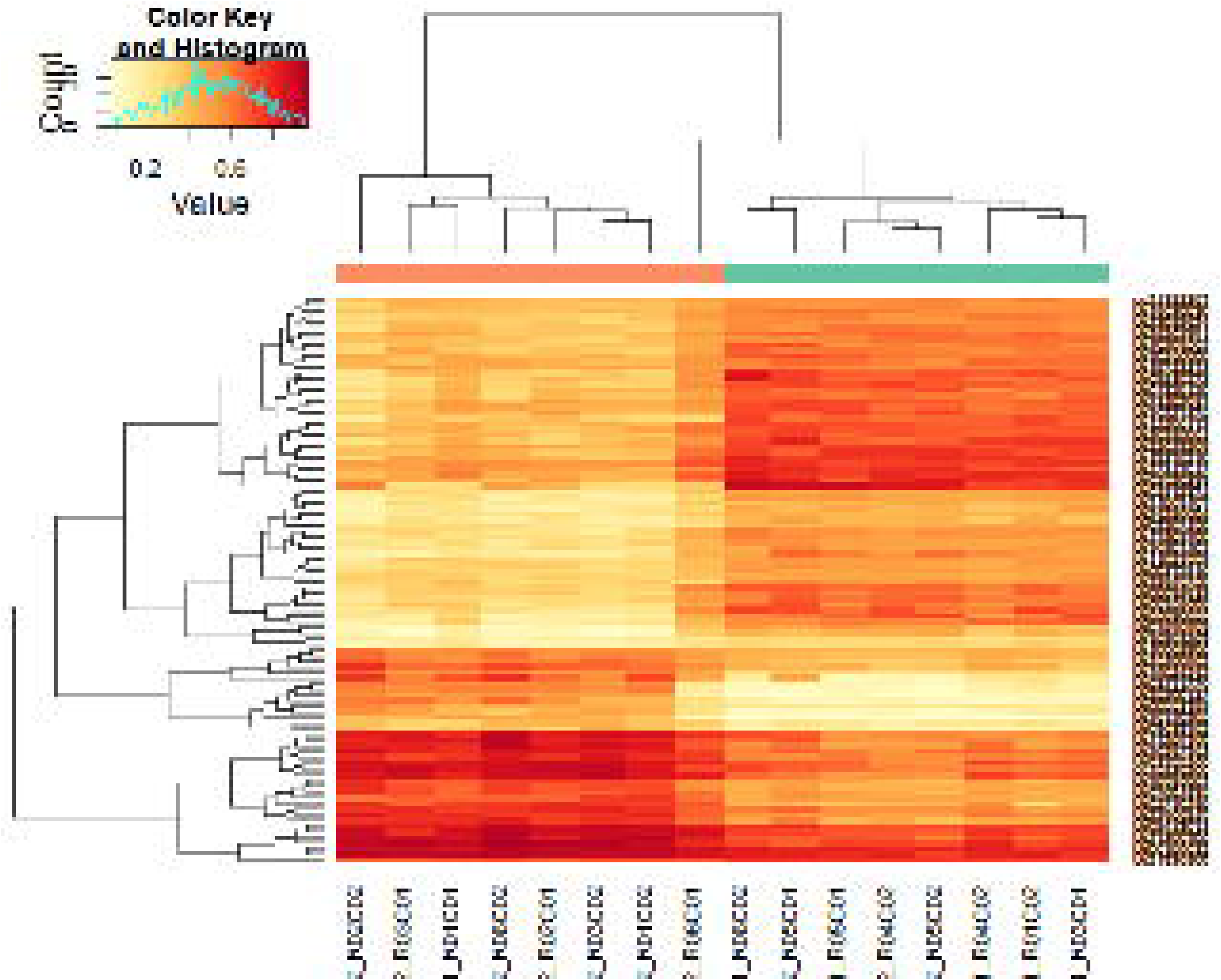
Hierarchical clustering of highly differentially methylated positions. Differentially methylated positions (DMPs) were sorted by F value (low intragroup variability and higher intergroup variability) and the top 250 sites were tested for clustering between study samples. Hierarchical clustering heatmap and dendogram are depicted in this figure, showing a clear separation of tumor samples from normal samples (top dendogram, control samples above green bar, tumor samples above orange bar). DMS median p-value heatmap shows a contrasting state of differential methylation between tumor and control samples indicating both gain and loss of differential methylation states in tumor tissues.

### Genomic Distribution of Differentially Methylated CpG sites

Differentially hypermethylated and hypomethylated sites displayed similar distribution with regard to gene elements as defined by HM450 -TSS1500, TSS200, First Exon, gene body, and 3UTR – **Fig 3.A**. However, they showed an asymmetric distribution with regard to CpG island relation with most of the hypermethylated sites mapping to CpG islands, whereas most of the hypomethylated sites mapped to open sea **Fig 3.B**.

**Figure 3:**
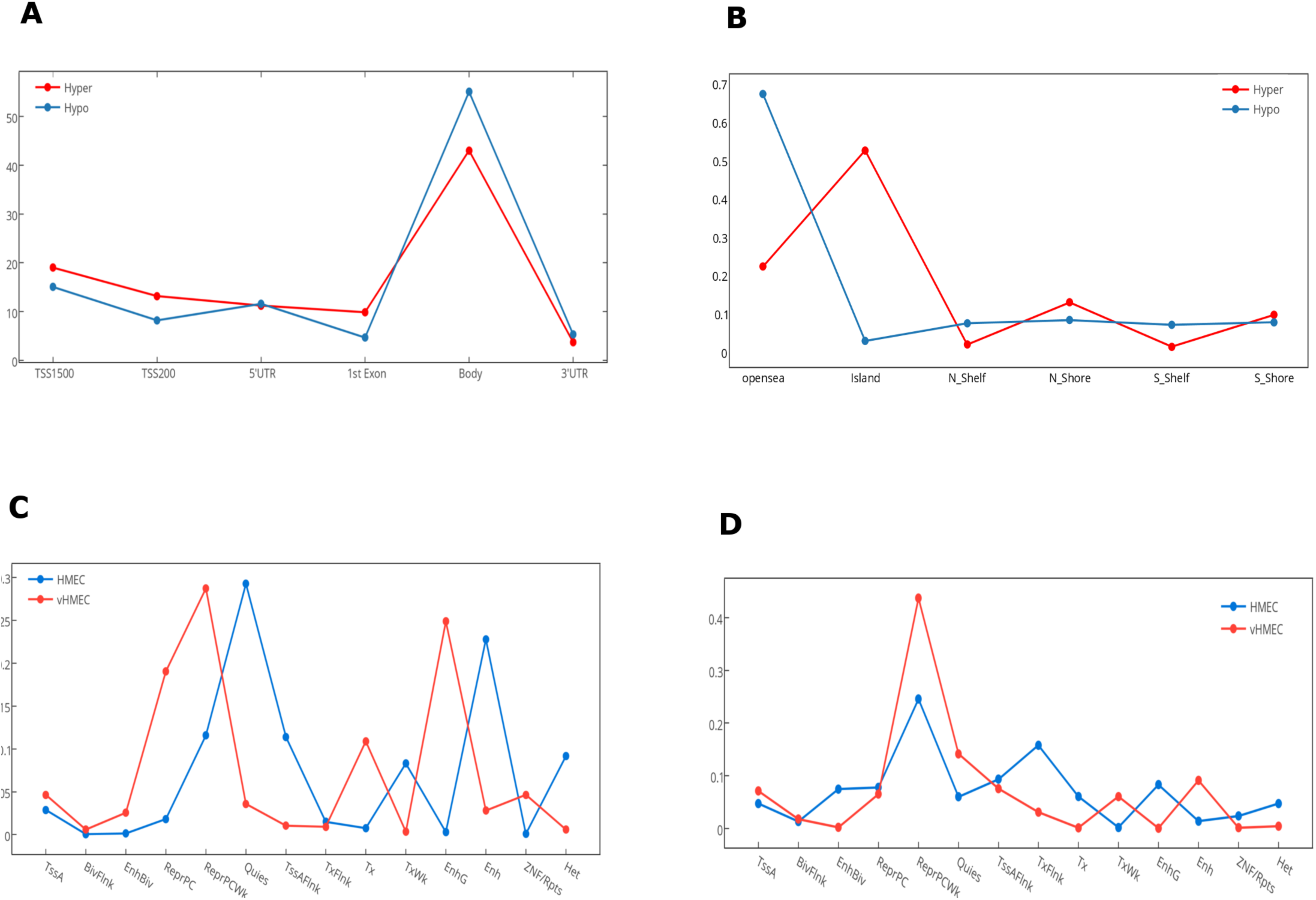
Genomic and epigenomic distribution of differentially methylated positions (DMPs). This figure details the number of DMPs in relation to gene elements, CpG islands and chromatin states. (A) Distribution of hyper and hypo methylated CpG sites in relation to gene elements. TSS: transcription start site, UTR: untranslated region. (B) Distribution of hyper and hypo methylated CpG sites in relation to CpG Islands. N_: north, S_: south (C) Distribution of Hypomethylated CpG sites in relation to chromatin states. (D) Distribution of Hypermethylated CpG sites in relation to chromatin states. Fourteen chromatin states are shown:

Of the 13633 hypermethylated sites, 24.37% (N=3323) mapped to Dnase hypersensitive sites compared with only 8.67% (N =568) of hypomethylated sites. Interestingly, while a greater percentage of hypermethylated compared to hypomethylated sites overlapped differentially methylated regions (DMR), [54.83% (N=1612), 11.47% (N=46)] respectively, hypomethylated sites were more concentrated in cancer DMR (CDMR), with 49.63% compared with 14.66% in hypermethylated sites, hypomethylated sites were more concentrated in cancer DMR (CDMR), with 49.63% compared with 14.66% in hypermethylated sites. The genomic distribution of hypermethylation and hypomethylation sites at each chromosome is shown in Additional File 1 and Additional File 2.

### Comparison to Reference Epigenome

We utilized data from the recently released Human epigenome reference data [19] to annotate the set of deferentially methylated CpG sites. We mapped hyper and hypo DMPs in the promoter region from our data against two reference epigenome breast cell lines: HMEC(Human mammary primary epithelial cells), and vHMEC (Human mammary primary epithelial cell variant)[20, 21]]. We examined the change in chromatin states – from the 15-chromatin states model [19] – that accompany the acquisition or loss of DNA methylation in the context of transitioning from normal to tumor states. Our results revealed a noticeable gain of repressive marks for the hypermethylated DMPs, which increased from 55.5% in HMEC cells to 78.7% in vHMEC cells. Interestingly, we also found a slight increase in the percentage of repressive marks in the hypomethylated DMPs, which increased from 54.3% to 61.6%. Notably, in both cases, most of the upsurge in repressive regions were concentrated in Polycomb-repressed regions **Fig 3.C, and 3.D**.

In addition, we observed a marked drop of all active chromatin states except for weak transcription and distal enhancer activity between the HMEC and vHMEC cells for the hypermethylated group. On the other hand, the hypomethylated group showed multiple notable shifts: From quiescent to Polycomb repression, from weak transcription to strong transcription, and from distal enhancers to genic enhancer (intronic enhancers).

### Candidate biomarkers discovery

Nimbl method was used for detection and prioritization of candidate biomarkers with greatest inter-group variability, and lowest intra-group variability[22]. Using this approach, we were able to identify a number of new as well as previously well-known BC biomarkers. Among the genes that showed significant promoter hypermethylation, we identified PAX6 [23, 24], WT1 [25], SOX1 [26], and TP73 [27, 28], all of them have been previously associated with BC. We also identified a set of previously uncharacterized biomarkers like PCDHGA1, HOXC4, and TBX15. To validate our candidate genes we interrogated our candidate gene list against BC methylome data from the Cancer Genome Atlas Network: http://cancergenome.nih.gov/ as compiled by MethHC [29] web portal. All the genes from our data were also significantly hypermethylated in the TCGA dataset. **Fig 4** shows promoter hypermethylation of the TP73 gene.

**Figure 4:**
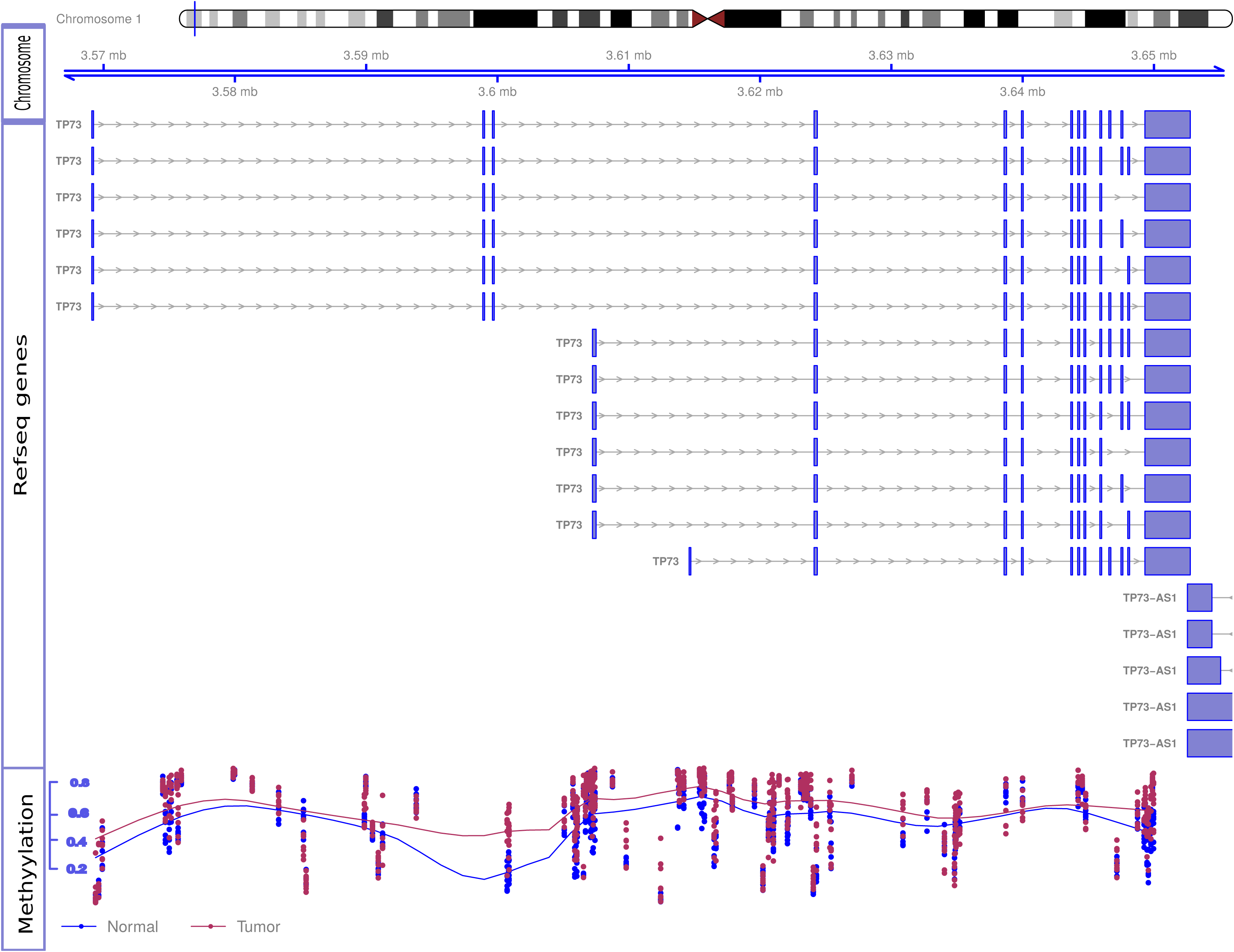
Hypermethylation of the TP73 gene. Differential methylation Beta-values for B tumor and B control samples at methylation array sites of TP73 gene are shown. The figure contains three tracks: the genomic location of the TP73 and its different RefSeq transcripts are shown in the ‘Chromosome’ and ‘RefSeq genes’ tracks respectively; the ‘Methylation’ track shows the methylation level in each tumor sample (red dot) and control sample (blue dot). The overall discordance in methylation Beta-values between tumor samples (red line in the methylation track) and control samples (blue line) is notable specially at TSS both for the long and short RefSeq transcripts (genomic areas around 3.56 mb and 3.6 mb, respectively). Tumor samples show relatively high beta-values compared to controls at these sites indicating differential promoter hypermethylation. TSS: Transcription Start Site.

### Pathway and Network Analysis

Results from the ReactomeFI for the EDG network uncovered a massive network of 1310 nodes (genes) and 5097 edges (interactions), while the EUG list produced a smaller network of 763 nodes and 2265 edges. Furthermore, loading the NCI (National Cancer Institute) cancer gene index identified 781, and 470, neoplasia related genes from the EDG, and EUG networks respectively, of which 332 EDG genes, and 222 EUG genes were associated with breast cancer in the cancer gene index.

Pathway enrichment analysis on the EUG network. Identified hippo signaling, Wnt signaling, and many extracellular matrix and metastasis promoting pathways as summarized in **Table 1**. Performing the pathway enrichment analysis on the breast cancer EUG subnetwork also identified hippo signaling and pathways of extracellular matrix in addition to pathways involved in immune response against viruses **Table 2**. Interestingly, breast cancer subnetwork showed significant enrichment for Epstein-Barr virus infection (FDR <0.001).

**Table 1:** Pathway enrichment analysis results for Epigenetically Upregulated Genes (EUG) interaction network. This pathway enrichment analysis and the interaction network were prepared using ReactomeFI Cytoscape app. The table shows the enriched pathways, the number of genes in the pathway from the total query gene set, and the number of genes in the pathway found in the interaction network. Results having <u>p-values <0.01</u> and a False Detection Rate < 0.001 are shown.

| Pathway | Number of genes in the Geneset | Number of genes in the Network | FDR |
| --- | --- | --- | --- |
| Hippo signaling pathway | 154 | 31 | <1.000e-03 |
| Arrhythmogenic right ventricular cardiomyopathy (ARVC) | 74 | 20 | <5.000e-04 |
| L1CAM interactions | 94 | 21 | 2.50E-04 |
| Wnt signaling pathway | 269 | 41 | 3.33E-04 |

**Table 2:** Pathway enrichment results for breast cancer related Epigenetically Upregulated Genes (EUG) subnetwork. ReactomeFI cytoscape app was used to extract breast cancer related subnetworks from EUG set by loading NCI cancer index and performing pathway enrichment analysis on interaction networks. Nodes that corresponded to malignant breast cancer were selected. The table shows the enriched pathways, the number of genes in the pathway from the total query gene set, and the number of genes in the pathway found in the interaction network. Results having p-values <0.01 and a False Detection Rate < 0.001 are shown.

| Pathway | Number of genes in the Geneset | Number of genes in the Network | FDR |
| --- | --- | --- | --- |
| CXCR4-mediated signaling events | 79 | 10 | 2.50E-04 |
| AP-1 transcription factor network | 70 | 9 | 7.27E-04 |
| HIF-1-alpha transcription factor network | 66 | 9 | 4.00E-04 |
| Viral myocarditis | 59 | 9 | 2.50E-04 |
| Pathways in cancer | 327 | 22 | <1.000e-03 |
| HTLV-I infection | 260 | 18 | 3.33E-04 |
| Proteoglycans in cancer | 225 | 17 | 2.00E-04 |
| Epstein-Barr virus infection | 202 | 16 | 1.67E-04 |
| Hippo signaling pathway | 154 | 14 | 3.33E-04 |
| Natural killer cell mediated cytotoxicity | 135 | 13 | 1.43E-04 |
| Alzheimer disease-presenilin pathway | 111 | 12 | 5.00E-04 |

Pathway analysis on the EDG network identified Neuroactive ligand-receptor interactions, G-protein signaling, RAP1 signaling, RAS signaling, Potassium channel signaling and many other pathways as summarized in **Table 3**. While the smaller EDG breast cancer subnetwork showed significant enrichment for a multitude of pathways including all the pathways that were enriched in the EDG network in addition to many cancer related and immune response pathways. Interestingly, the EDG sub network was also significant for direct p53 effectors. The complete list of enriched pathways for the EDG breast cancer subnetwork is shown in **Additional file 3: Table S1**.

**Table 3:** Pathway analysis on the Epigenetically Downregulated Genes (EDG) interaction network. The functional interaction network was constructed using ReactomeFI cytoscape app. The table shows the enriched pathways, the number of genes in the pathway from the total query gene set, and the number of genes in the pathway found in the interaction network. Results having p-values <0.01 and a False Detection Rate < 0.001 are shown.

| Pathway | Number of genes in the Geneset | Number of genes in the Network | FDR |
| --- | --- | --- | --- |
| Neuroactive ligand-receptor interaction | 275 | 81 | <1.000e-03 |
| GPCR ligand binding | 433 | 107 | <5.000e-04 |
| PI3K-Akt signaling pathway | 346 | 81 | <3.333e-04 |
| Extracellular matrix organization | 263 | 65 | <2.500e-04 |
| Pathways in cancer | 327 | 74 | <2.000e-04 |
| Rap1 signaling pathway | 213 | 54 | <1.667e-04 |
| Regulation of actin cytoskeleton | 215 | 54 | <1.429e-04 |
| Neurotransmitter Receptor Binding And Downstream Transmission In The Postsynaptic Cell | 137 | 40 | <1.250e-04 |
| Potassium Channels | 86 | 30 | <1.111e-04 |
| Heterotrimeric G-protein signaling pathway-Gi alpha and Gs alpha mediated pathway | 147 | 41 | <1.000e-04 |
| Proteoglycans in cancer | 225 | 54 | <9.091e-05 |
| Ras signaling pathway | 227 | 54 | <8.333e-05 |
| ECM-receptor interaction | 86 | 28 | 1.54E-04 |
| Calcium signaling pathway | 181 | 45 | 2.14E-04 |
| FGF signaling pathway | 92 | 29 | 2.00E-04 |
| Focal adhesion | 206 | 48 | 2.50E-04 |
| Gastrin-CREB signalling pathway via PKC and MAPK | 207 | 48 | 2.35E-04 |
| Cell adhesion molecules (CAMs) | 143 | 37 | 2.78E-04 |
| Wnt signaling pathway | 269 | 57 | 4.21E-04 |
| MAPK signaling pathway | 259 | 55 | 4.00E-04 |
| Heterotrimeric G-protein signaling pathway-Gq alpha and Go alpha mediated pathway | 108 | 30 | 5.24E-04 |
| IL4-mediated signaling events | 63 | 21 | 7.73E-04 |
| HTLV-I infection | 260 | 54 | 7.83E-04 |
| GABAergic synapse | 90 | 26 | 7.92E-04 |
| Signaling by Type 1 Insulin-like Growth Factor 1 Receptor (IGF1R) | 86 | 25 | 8.40E-04 |
| Retrograde endocannabinoid signaling | 103 | 28 | 8.85E-04 |
| Melanoma | 71 | 22 | 9.26E-04 |

## Discussion

### Reference Human Epigenome Annotations

The recent release of the human reference epigenome data by the Roadmap project ushered in a new era of epigenomics. The current study utilized this new wealth of information to interpret epigenetic data. We successfully mapped hyper and hypo DMPs to chromatin states from normal and premalignant reference breast cells (HMEC and vHMEC respectively). Despite the fact that vHMEC is a premalignant and not a primary tumor cell, we argue that vHMEC is a suitable model for the epigenetic changes that accompany BC tumorigenesis because the vast majority of epigenetic changes tend to occur early during BC tumorigenesis [30–33].

Notably, our data revealed a strong Polycomb repression in both hypermethylated and hypomethylated CpG sites. These findings, are in accordance with the emerging evidence that DMPs are enriched for Polycomb repression in primary breast tumors [34] and triple negative BC [35]. Moreover, Various elements of the Polycomb repressive complexes are well known to be overexpressed in BC [36, 37] and are required for stem cell state in mammary tumors [38, 39]. Interestingly, Reyngold et al, found that unlike primary tumors, genes methylated in metastatic lesions seem to lack Polycomb repressive marks [40].

### Network-Based pathway enrichment analysis

Our network-based pathway enrichment results for the EUG network revealed many upregulated pathways that have been previously associated with BC tumorigenesis. Hippo signaling, which appeared as the top significantly enriched pathway in our results, has recently emerged as an important regulator of BC growth, migration, invasiveness, stemness, as well as drug resistance[41]. Wang et al, demonstrated that overexpression of YAP enhanced BC formation and growth. Hiemer et al, found that both TAZ and YAP -key effectors of the Hippo pathway -are crucial to promote and maintain TGFβ-induced tumorigenic phenotypes in breast cancer cells [42]. In addition, YAP was demonstrated to mediate drug resistance to RAF and MEK targeted cancer therapy [43, 44]. Interestingly, we also reported an upregulated Wnt signaling pathway, which has been linked to BC growth and malignant behavior [45]. Jinhua et al found that Wnt signaling pathway is required for triple-negative breast cancer development [46]. Recent studies have suggested long lasting reduced Wnt signaling as the mechanism by which early pregnancy protects against BC [47].

Regarding the EDG network, Neuroactive ligand-receptor interaction, in addition to GPCR, RAS and Rap1 signaling were among the most significantly enriched pathways. Recent studies have found Neuroactive ligand-receptor interaction related genes to be hypermethylated in colorectal and EBV associated gastric cancers [20,21]. Elements of RAS signaling like RASSF has been frequently found to be hypermethylated in BC [49], moreover, Qin et al, has demonstrated that resveratrol is able to demethylate RASSF1 promoter through decreased DNMT1 and DNMT3b in mammary tumors [50, 51]. Notably, we reported the apparent silencing of multiple pro-tumor pathways in our results like GPCR and RAP1 signaling, the precise significance of this findings remains unclear. In addition, we also noticed the bivalent enrichment of multiple pathways (where different elements of the same pathway are both up and down regulated). Interpreting such perturbations is tricky, and predicting the net outcome of those perturbations might not be readily obvious given the crosstalk between different pathways.

### EBV signature

We previously reported a strong association between EBV and BC in Sudanese patients [52], we also reported frequent epigenetic silencing of major tumor suppressor genes coupled with low frequency of known tumor associated mutations in the same population [53]. In this study, we have demonstrated genome-wide epigenetic alterations consistent with our original proposition that epigenetic changes are the primary driver of BC tumorigenesis in Sudanese patients.

A myriad of recent studies point toward a common theme in EBV associated cancers characterized by genome-wide epigenetic changes coupled with a paucity of mutations. EBV infection is now known to play significant role in epithelial cancers like nasopharyngeal and gastric carcinomas mainly through genome-wide epigenetic changes [54–56]. Li et al observed a unique epiphenotype of EBV associated carcinomas suggesting a predominant role for EBV infection in the ensuing epigenetic dysregulation of those cancers [17]. Another study attributed the genome-wide promoter methylation in EBV driven gastric cancer to the induced expression of DNA methyltransferase-3b (DNMT3b) [57].

Our data mirrored the overall unique pattern of EBV infection characterized by sweeping epigenetic changes accompanied by low mutation frequency. Furthermore, we also showed that the EUG network was significantly enriched for EBV infection pathway **Fig 5**. In addition, results from MSig perturbations obtained from GREAT web tool (which predicts functions of cis regulatory elements) [58], showed significant enrichment for a set of downregulated genes which had been previously correlated with increased expression of EBV EBNA1 protein in NPC, in the hypermethylated CpG sites group, data not shown.. For the hypomethylated CpG group, we found genes upregulated in B2264–19/3 cells (primary B lymphocytes) within 30–60 min after activation of LMP1 to be significantly enriched in MSig oncogenic signature. These findings taken together provide the first bioinformatics evidence of a possible active role for EBV infection in BC tumorigenesis in Sudanese patients.

**Figure 5:**
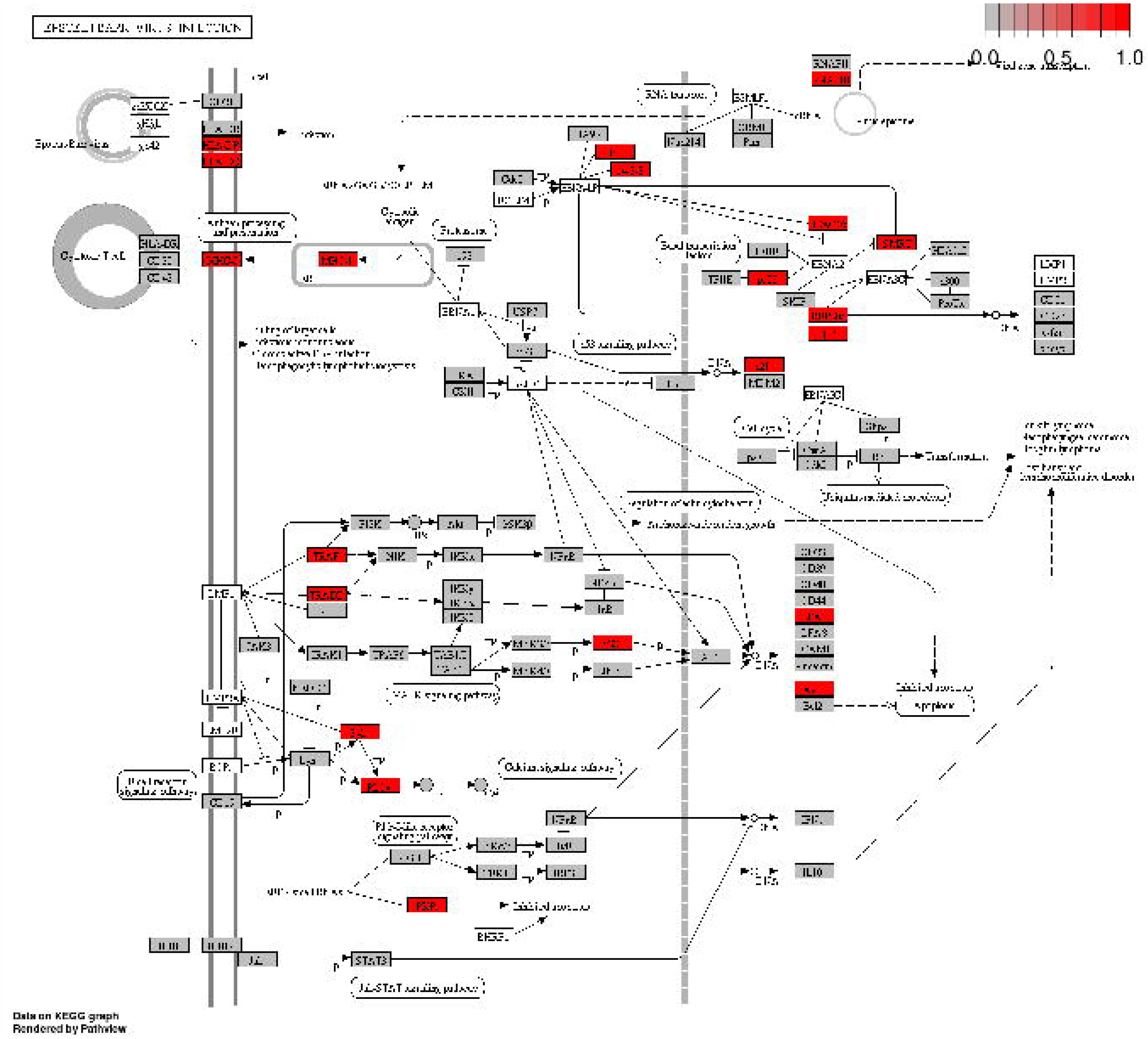
Tumor Epigenetically Upregulated Genes (EUG) in Epstein-Barr Virus Infection pathway. Many genes bearing methylation marks that promote gene expression (hypomethylation in the promoter area and first exon or hypermethylation in the gene body region) – referred to in this study Epigenetically Upregulated Genes – were found to be integral parts of EBV Infection KEGG pathway (highlighted red and grey boxes). This group of genes showed significant enrichment for Epstein-Barr Virus Infection Pathway (red boxes are highly enriched nodes). Epstein-Barr Virus Infection KEGG Pathway was obtained from KEGG pathways database [http://www.kegg.jp/pathway/hsa05169].

## Conclusions

In conclusion, our study uncovered interesting epigenetic patterns characterized by increased acquisition of Polycomb repressive marks, as revealed from comparison to human reference epigenome breast cells. We identified many potential BC biomarkers like TP73, and TBX15. We also identified many significantly enriched developmental pathways including Hippo and Wnt signaling pathways. Additionally, our bioinformatics analysis indicated a possible role for EBV infection in BC tumorigenesis.

## Materials and Methods

### Ethical considerations

Ethical approval for this study was obtained from the Institute of Endemic Diseases, University of Khartoum Ethical Committee. Written informed consent was obtained from all participants; all clinical investigations were conducted according to the principles expressed in the Declaration of Helsinki: http://www.wma.net/en/30publications/10policies/b3/index.html

Samples, DNA extraction, and Illumina Infinium HumanMethylation 450 (HM450) array Genomic DNA was extracted from eight samples of primary breast tumors and eight normal samples from adjacent healthy. All samples were independently reviewed by histopathologists. DNA was extracted from tissues using Promega genomic DNA purification kit [59] following the standard protocol as described by the manufacturer. DNA methylome profiling was performed using Illumina Infinium HumanMethylation 450 (HM450) [60]BeadChip array by Beijing Genomics Institute (BGI)

### Data Preprocessing

For quality control, any array probes with p detection value less than 0.05 or missing beta values were removed. In addition, array sites corresponding to sex chromosomes or mapping to SNPs were filtered out. Peak-based correction[61] (PBC) was used to normalize the final dataset and to correct for probe type bias. Density plots of beta values for individual samples are shown in Additional File 4**: Fig S3**

### Genome-wide DNA Differential methylation Analysis

A trilateral approach consisting of two statistical methods augmented by one numerical method was used for the differential methylation analysis: Moderated T test from R limma [62] package; Wilcoxon test (Non Parametric test) from R stat package; and Nimbl [22] (Numerical Identification of Methylation Biomarker Lists) which is a Matalab package designed to identify and prioritize differentially methylated sites.

Nimbl core module identify potential biomarkers by calculating a score based on the inter-group and intra-group variability:

Score = beta_valdist – (mediandiff – beta_valdist)

Where beta_valdist is the distance in beta values between non-overlapping groups and mediandiff is the absolute difference of the medians of each group [22]. It then assigns high scores for CpG sites that achieve higher discrimination between groups while maintaining low within-group variability. Nimbl-compare module was also used to extract the final set of CpG sites that were identified by all three methods. Hierarchical clustering analysis was performed using the top 250 differentially methylated sites sorted by F value.

### Reference Epigenome Annotations

Bed files of chromatin states for both HMEC and vHMEC cells were obtained from Roadmap web portal: http://egg2.wustl.edu/roadmap/web_portal/, further analysis was performed in GALAXY web-based platform [63–65] and R statistical software.

### Network and Pathway Analysis

Differential methylation analysis produced two lists of differentially methylated genes (hyper and hypo) and their enrichment of differentially methylated sites in their gene regions, i.e., promoter region, gene body, 3UTR, etc. The aggregated gene list was sorted by the count of methylated sites in the promoter area, first exon, and gene body regions. Subsequently all epigenetically upregulated genes (EUG) were combined in a single group, i.e., genes bearing methylation marks that promote gene expression – hypomethylation in the promoter area, and first exon or hypermethylation in the gene body region- in a single group. Then we compiled a second group of epigenetically downregulated genes (EDG), i.e., genes bearing methylation marks that inhibits gene expression, i.e., hypermethylation of the promoter area, and the first exon or hypomethylation of the gene body region. We excluded other gene-based regions that are not well correlated with gene expression from further analysis.

We utilized ReactomeFI[66], a Cytoscape [67] app to perform network and pathways analysis. Projecting the lists of EDG and EUG groups through the ReactomeFI functional network produced two corresponding networks. To extract breast cancer specific subnetworks from EUG and EDG groups we loaded NCI cancer index from within the ReactomeFI app, and we selected nodes that corresponded to malignant breast cancer.

**Abbreviations**
BC: breast cancer
DMP: differentially methylated position
DMR: differentially methylated region
CDMR: cancer differentially methylated site
TSS: transcription start site
UTR: untranslated region
MSig: mutation signature

## Authors’ contributions

HSM conceived and design the study and contributed to manuscript writing and data interpretation. MEI contributed to study design and manuscript writing. MOA performed the data analysis, contributed to interpretation and prepared the manuscript draft. JHS and AHF recruited patients and provided samples. MK contributed to data analysis. AME performed the histopathology, UKA, MAS, RA, and GS contributed to sample collection, DNA extraction and purification. All authors read and approved the final manuscript

## Acknowledgments

We thank the breast cancer patients for their participation in this study. This work is dedicated to deceased Mohammed Abdelrazig. Senior Surgeon at Khartoum teaching hospital who facilitated samples collection.

## Additional Files

**Additional File 1 (Fig S1):** Genomic distribution of hypermethylation marks shown at each chromosome. Black color indicates hypermethylation sites. File format: PDF.

**Additional File 2 (Fig S2):** Genomic distribution of hypomethylation marks shown at each chromosome. Black color indicates hypermethylation sites. File format: PDF.

**Additional File 3 (Table S1):** Pathway enrichment results for breast cancer related Epigenetically Downregulated Genes (EDG) subnetwork. ReactomeFI cytoscape app was used to extract breast cancer related subnetworks from EUG set by loading NCI cancer index and performing pathway enrichment analysis on interaction networks. Nodes that corresponded to malignant breast cancer were selected. The table shows the enriched pathways, the number of genes in the pathway from the total query gene set, and the number of genes in the pathway found in the interaction network. Results having p-values <0.01 and a False Detection Rate < 0.001 are shown. File format: DOCX.

**Additional File 4 (Fig S3):** Density plots of beta values for individual samples. Shades of red and yellow colors represent tumor samples, whereas shades of blue and green represent normal samples. File format: PDF.

